# Forest change within and outside protected areas in the Dominican Republic, 2000-2016

**DOI:** 10.1101/558346

**Authors:** John D. Lloyd, Yolanda M. León

## Abstract

We used Landsat-based estimates of tree cover change to document the loss and gain of forest in the Dominican Republic between 2000 and 2016. Overall, 2,795 km^2^ of forest were lost, with forest gain occurring on only 393 km^2^, yielding a net loss of 2,402 km^2^ of forest, a decline of 11.1% or 0.7% per year. Deforestation occurred in all of the major forest types in the country, and ranged from a 13% decline in the area of semi-moist broadleaf forest to a 5.9% loss of cloud forest, mostly attributed to agriculture. Fire was a significant driver of forest loss only in Hispaniolan pine (*Pinus occidentalis*) forests and, to a lesser extent, in adjacent cloud forest. Deforestation rates were lower within protected areas, especially in dry and semi-moist broadleaf forests at lower elevations. Protected areas had a smaller, and generally negligible, effect on rates of forest loss in pine forest and cloud forest, largely due to the effects of several large wildfires. Overall, rates of deforestation in the Dominican Republic were higher than regional averages from across the Neotropics and appeared to have accelerated during the later years of our study period. Stemming deforestation will likely require enforcement of prohibitions on large-scale agricultural production within protected areas and development of alternatives to short-cycle, shifting agriculture.

## Introduction

Human well-being is linked inextricably with the fate of the planet’s forests. Forests provide goods and income to the rural poor throughout the developing world [1], generate employment for more than 10 million people throughout the world [2], yield renewable flows of raw materials for commercial and domestic use, sustain stable flows of clean water [3,4], buffer against local extremes of climate [5], and regulate global climate and carbon cycles [6,7]. Indeed, the very persistence of modern human societies may be incompatible with the conditions created by ongoing deforestation [8]. The survival of an uncounted number of non-human species also depends on the persistence of forested landscapes.

Efforts to conserve Earth’s remaining forests, and to understand the consequences of their disappearance, demand estimates of where, and at what rate, forest loss is occurring [9,10]. Reliable national-level data on forests is urgently needed to inform policies on forest conservation, sustainable development, and climate-change mitigation. A significant contribution to these efforts was made by Hansen et al. [9], who provided satellite-based estimates of global forest cover at a relatively fine temporal and spatial scale. Those data have been used subsequently to generate regional estimates of deforestation [11], estimates of loss of specific forest types [12], and country-specific descriptions of forest change [13]. Although analyses at planetary and regional scales provide useful insights for efforts to limit the deleterious consequences of global change [14] or meet global sustainable development goals [15], smaller-scale analyses, especially at the national or sub-national level, are useful because they align more closely with the level at which policies on forest use and conservation are implemented. Thus, country-specific analyses of deforestation allow for the evaluation of the efficacy of conservation interventions and, ideally, implementation of adaptive changes as needed.

Here, we examine spatial and temporal patterns of change in forest cover in the Dominican Republic (DR) between 2000 and 2016 using Hansen et al.’s [9] forest-cover dataset and its annual updates. In particular, we document changes in the extent of forest cover, by forest type, and examine the efficacy of the nation’s system of protected areas - the country’s primary conservation tool - in stemming forest loss. We focused on the DR for several reasons. First, as a middle-income country, it is broadly reflective of the changing dynamics and challenges faced globally in conserving forests in developing countries experiencing rapid economic growth: the DR’s average economic growth of 5.3% over the past 25 years has been among the strongest in Latin America and the Caribbean [16]. Second, it supports an outstanding number of forest-dependent endemic plants and animals [17,18], many of which are threatened with extinction [19]. Third, very little published, quantitative information exists on the status of forests in the DR. Only two studies have produced quantitative estimates of change in forest cover [20,21], and none that we are aware of have produced estimates specific to the different forest types in the country. In quantifying recent changes in the extent of different forest ecosystems in the DR, we hope to provide an initial evaluation of forest-specific conservation policies, identify spatial hotspots of deforestation and forest types at greatest risk, and to suggest fruitful areas for investment of conservation resources.

## Methods

### Quantifying forest change

We estimated annual changes in forest cover in the DR between 2000 and 2016 using version 1.4 of the Hansen et al. [9] tree cover data, accessed through the Google Earth Engine [22]. The baseline year for these data is 2000, at which time percent tree cover was estimated in every 30-m pixel. Tree cover was defined by Hansen et al. [9] as all vegetation >5 m tall. In each subsequent year, every pixel can either remain in a forested state or undergo deforestation, which is defined as the transition to an entirely unforested state at the Landsat pixel level. Partial removal of forest canopy is not considered loss in the scope of this analysis. Forest gain, conversely, is defined as the transition from unforested to >50% tree cover during the period 2000-2012; forest gain is not calculated on an annual basis nor does it include regrowth after 2012. We used the per-pixel estimate of tree cover in 2000 as our baseline such that our estimates of the area of forest cover lost or gained are corrected for initial conditions. For example, a pixel (900 m^2^) that was estimated to have had 25% tree cover in 2000, and that was identified as having been deforested between 2000 and 2016, was calculated to have contributed a loss of 225 m^2^ of forest (i.e., total pixel area multiplied by the percent of forest cover in 2000).

We calculated change in the extent of forest in two ways. First, we estimated change from 2000-2016 for all areas identified as forested in 2000 by Hansen et al. [9]. This provides a broad overview of changes in tree cover across the country, including not only in natural forest but also in heavily managed areas like forested parks in urban areas or agroforestry plantations. To gain insight into patterns of change in naturally forested areas, we also generated separate estimates of change for each major forest type identified in the 1996 land-cover map of Tolentino and Peña [23], which provides the only pre-2000 estimate of land-cover types across the country. For this portion of the analysis, we focused only on natural, unmanaged forests, thus excluding urban parks, tree plantations (e.g., mango [*Mangifera* spp.], coconut [*Cocos nucifera*], and oil palm [*Elaeis guineensis*]), and shade crops (e.g., coffee [Coffea arabica] or cacao [*Theobroma cacao*]).

The forest types considered in the second analysis include Hispaniolan pine (*Pinus occidentalis*) forest, which was classified by Tolentino and Peña [23] into both an open (“bosque conífera abierto”) and closed-canopy (“bosque conífera denso”) category; cloud forest (“bosque nublado”); moist broadleaf forest (“bosque húmedo”); semi-moist broadleaf forest (“bosque semihúmedo”); and dry forest (“bosque seco”). Pine forests occur at the highest, coldest elevations of the Sierra de Bahoruco, Sierra de Neiba, and Cordillera Central. Cloud forest usually arises at the lower elevational limit of pine, in areas with mild temperatures, abundant precipitation, and persistent ground-level clouds. Cloud forest transitions into moist broadleaf forest at lower elevations where average annual precipitation drops below ∼2,000 mm [23]. Moist broadleaf forest occupies a broad elevational range, from near sea level in wet areas in the north of the country, such as Los Haitises, to ∼1800 m on drier mountain slopes, such as the southern slope of Sierra de Bahoruco. Semi-moist broadleaf forest occurs in coastal areas and on mountain slopes as a transitional zone between dry forest and moist broadleaf forest. Dry forest is the only non-evergreen forest, found at relatively low elevations (< 500 m) with a warm, dry, and seasonal climate. Unlike the other forest types, most extant dry forest is secondary forest in the process of recovering from past anthropogenic disturbances [23].

### Separating wildfire from other causes of forest loss

Fire can be a significant driver of vegetation dynamics in the DR, especially in montane forests [24], so to examine the role of wildfire as an agent of forest loss we used the monthly, MODIS-based estimates of the global area burned [25]. We aggregated monthly estimates of area burned for each year and assumed that forest loss was caused by fire for any pixel in the Hansen et al. [9] data that was within the boundaries of a burned area and was estimated as having been deforested in that year.

### Quantifying forest change within protected areas

The DR has an extensive national protected area system, covering 26% of its territory [26]. To examine whether forest within formally protected areas showed different patterns of change, we calculated forest change and area burned for each protected area within the DR.

## Results

### Quantifying forest change

Trees covered 21,494 km^2^ of the DR in 2000, roughly 45% of its total land area. Deforestation removed 2,795 km^2^ of this tree cover by 2016, while reforestation or afforestation occurred on only 393 km^2^, a net loss of 2,402 km^2^, reducing forest cover to roughly 40% of the territory. This amounts to an 11.1% decline in forest cover at the national level over the period of analysis, an annual deforestation rate of 0.7%.

Considering only the DR’s major natural forest types, forest cover shrank from 9,517 km^2^ in 2000 to 8,644 km^2^ in 2016, a net loss of 9.2% (Table 1). Depending on forest type, this change ranged from −5.9% in cloud forests to −13.1% in semi-moist forests. The extent of loss varied among years but, with the exception of dry forest, tended to increase after 2010 (Fig 1). Forest gain was negligible in all of the natural forest types.

**Table 1.**
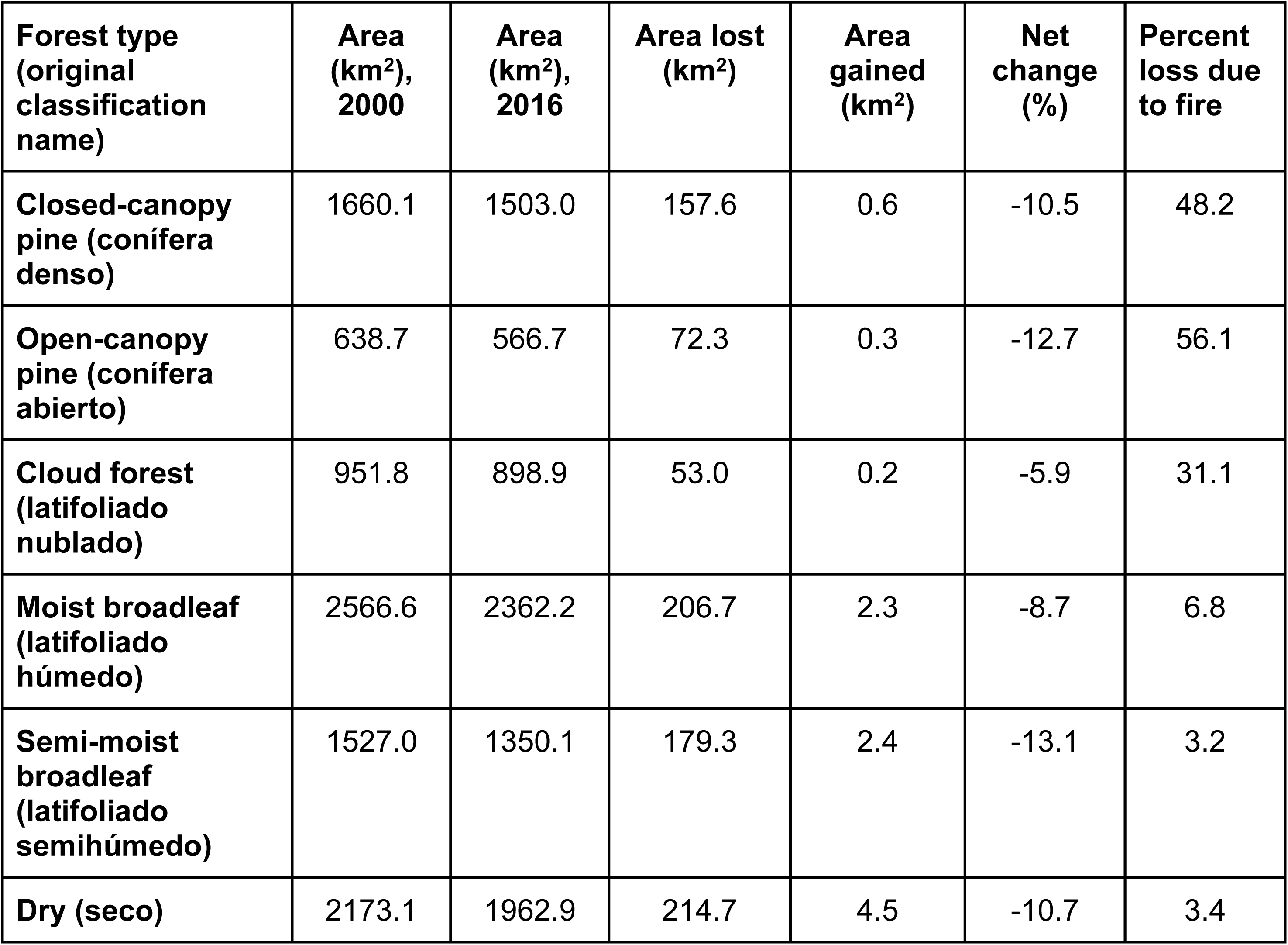
Changes in the estimated areal extent of forest in the Dominican Republic between 2000 and 2016 for the 6 forested land-cover types identified in national land-cover mapping.

| Forest type<br>(original<br>classification<br>name) | Area<br>(km<sup>2</sup>),<br>2000 | Area<br>(km<sup>2</sup>),<br>2016 | Area lost<br>(km<sup>2</sup>) | Area<br>gained<br>(km<sup>2</sup>) | Net<br>change<br>(%) | Percent<br>loss due<br>to fire |
| --- | --- | --- | --- | --- | --- | --- |
| Closed-canopy<br>pine (conífera<br>denso) | 1660.1 | 1503.0 | 157.6 | 0.6 | -10.5 | 48.2 |
| Open-canopy<br>pine (conífera<br>abierto) | 638.7 | 566.7 | 72.3 | 0.3 | -12.7 | 56.1 |
| Cloud forest<br>(latifoliado<br>nublado) | 951.8 | 898.9 | 53.0 | 0.2 | -5.9 | 31.1 |
| Moist broadleaf<br>(latifoliado<br>húmedo) | 2566.6 | 2362.2 | 206.7 | 2.3 | -8.7 | 6.8 |
| Semi-moist<br>broadleaf<br>(latifoliado<br>semihúmedo) | 1527.0 | 1350.1 | 179.3 | 2.4 | -13.1 | 3.2 |
| Dry (seco) | 2173.1 | 1962.9 | 214.7 | 4.5 | -10.7 | 3.4 |

**Fig 1.**
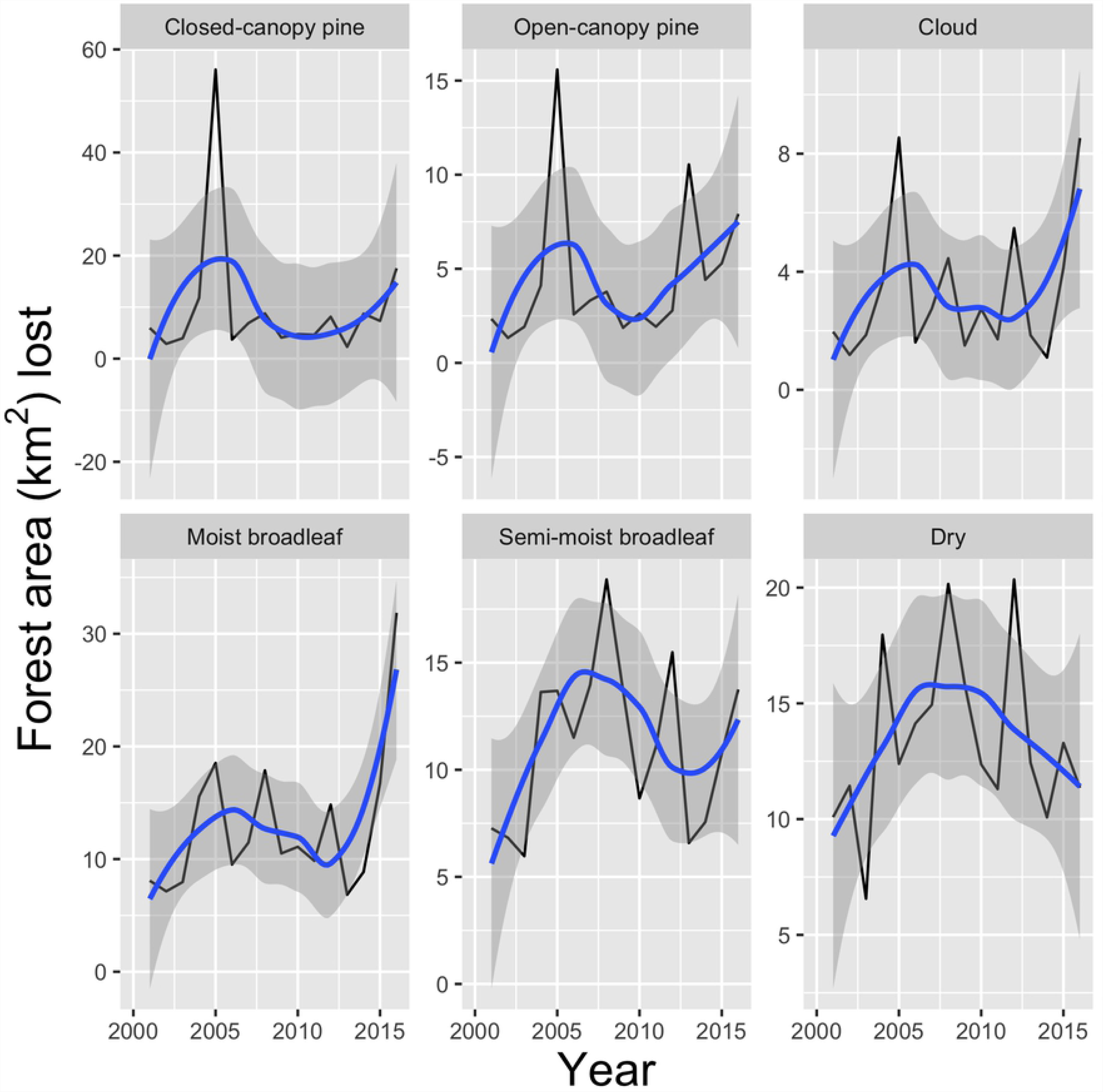
Deforestation rates in the Dominican Republic from 2000-2016 in each of the country’s major upland forest types. The extent of deforestation (solid black line) varied among years and among the six major forest types. When smoothed via loess (solid blue line; shaded interval is 95% confidence interval), deforestation appeared to accelerate after 2010, with the exception of dry forest.

### Fire as a deforestation driver

Fire accounted for a significant amount of loss in the Hispaniolan pine forests, in both open- and closed-canopy types (Table 1). However, most of the loss caused by fire occurred in a single year (Fig 2). In closed-canopy pine forest, fires in 2005 accounted for roughly 77% of the area burned between 2000 and 2016; in open-canopy pine, 50%; in cloud forest, 53%; and in moist broadleaf, 55%. Lesser peaks occurred in 2014 for both pine forest types (12% and 6% of total area burned over the course of the study for closed-canopy and open-canopy, respectively) and again in 2015 with losses for closed-canopy pine (4%), open-canopy pine (10%), cloud forest (19%), and moist broadleaf (32%). Fire accounted for relatively little loss in area among the dry and semi-moist broadleaf forest types.

**Fig 2.**
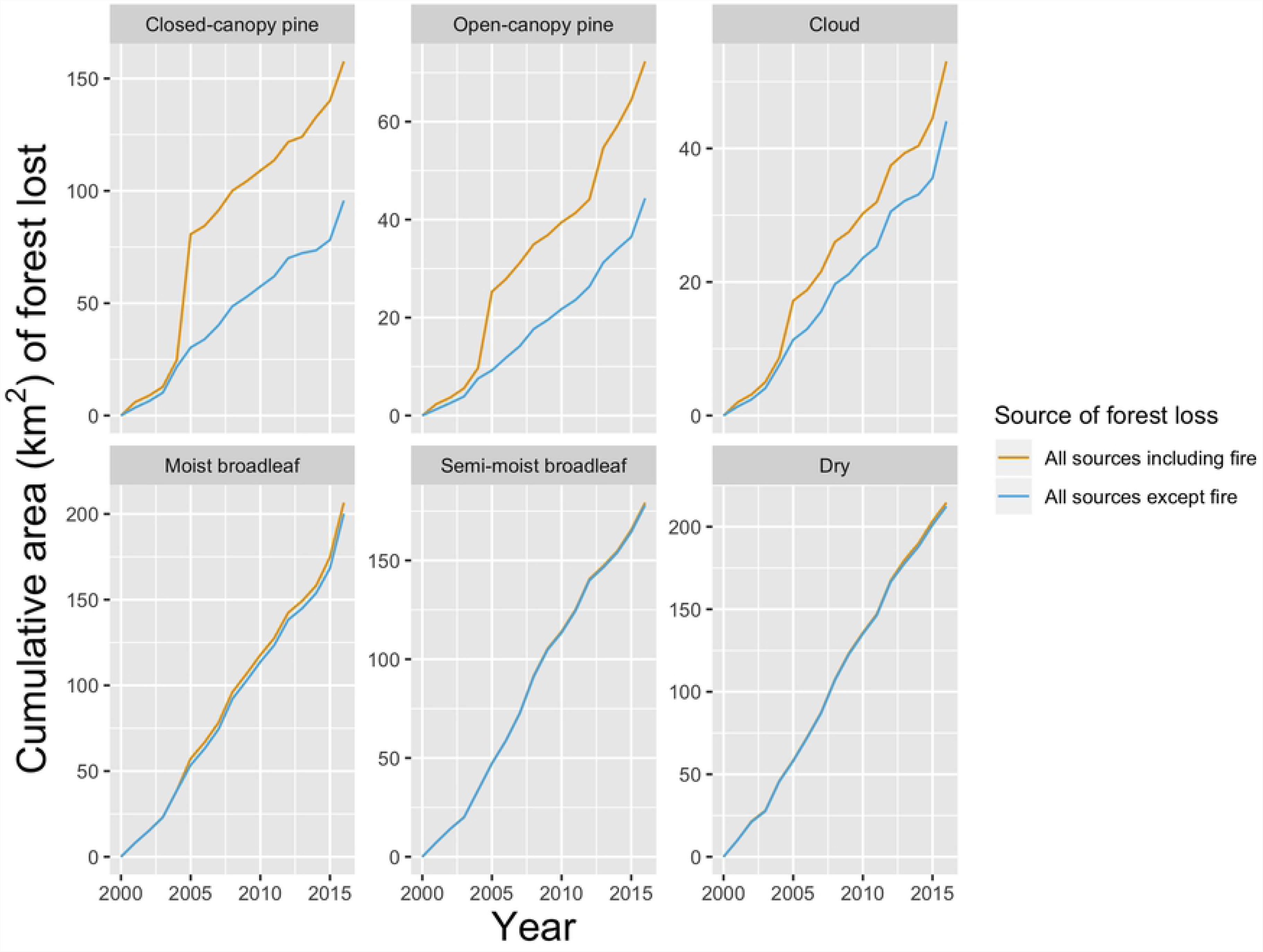
Relative importance of fire and other sources as agents of forest loss in major upland forest types of the Dominican Republic, 2000-2016. Declines in the extent of the six major forest types in the Dominican Republic were driven in most years by forest loss from sources other than fire (blue lines); significant losses due to fire were apparent only in 2005, as shown by the gap between the amount of forest lost to all sources (orange lines) and the amount of forest lost to sources other than fire. Lesser peaks in area burned were apparent in 2015 for both pine types and cloud forest.

### Protected area deforestation

Rates of deforestation within protected areas largely mirrored overall trends in forest change, except for dry and semi-moist broadleaf forests where forest loss was substantially lower within protected areas (Table 2). Within protected areas, forest accounted for 7,381 km^2^ in 2000, covering 57% of the land. By 2016, forest cover had shrunk by 670 km^2^ (−8.5%) and covered only 52% of the land in protected areas.

**Table 2.** Changes in the estimated areal extent of forest within formally protected areas in the Dominican Republic between 2000 and 2016 for the 6 forested land-cover types identified in national land-cover mapping.

| Forest type<br>(original<br>classification<br>name) | Percent<br>protected | Area<br>(km<sup>2</sup>),<br>2000 | Area lost<br>(km<sup>2</sup>) | Area<br>gained<br>(km<sup>2</sup>) | Net<br>change<br>(%) | Percent<br>loss<br>due to<br>fire |
| --- | --- | --- | --- | --- | --- | --- |
| Closed-canopy pine<br>(conifero denso) | 81.6 | 1354.2 | 126.2 | 0.4 | -10.3 | 47.3 |
| Open-canopy pine<br>(conifero abierto) | 56.9 | 363.1 | 49.4 | 0.2 | -15.7 | 54.7 |
| Cloud forest<br>(latifoliado nublado) | 78.1 | 743.6 | 39.2 | 0.2 | -5.5 | 20.9 |
| Moist broadleaf<br>(latifoliado humedo) | 43.8 | 1122.4 | 73.1 | 0.5 | -6.9 | 7.7 |
| Semi-moist<br>broadleaf (latifoliado<br>semi-humedo) | 40.0 | 610.4 | 43.3 | 0.5 | -7.5 | 1.2 |
| Dry (seco) | 49.1 | 1066.8 | 43.6 | 0.3 | -4.2 | 1.8 |

Protected areas offered little defense against fire, either. Fire accounted for significant amounts of the estimated loss of both pine and cloud forests within protected areas (Table 2). In fact, of the three forest types experiencing significant losses due to fire, nearly all of the burned area occurred within protected areas: for closed-canopy pine forest, 96% of the burned area was within a protected area; for open-canopy pine forest, 91%; and for cloud forest, 91%.

Extent of forest loss varied substantially among protected areas for all forest types. The largest losses in pine forests occurred in José del Carmen Ramírez National Park (JC Ramírez), largely due to the large 2005 fire, followed by Sierra de Bahoruco National Park (Bahoruco) due to sources other than fire (Figs. 3 and 4; S1 File).

**Fig 3.**
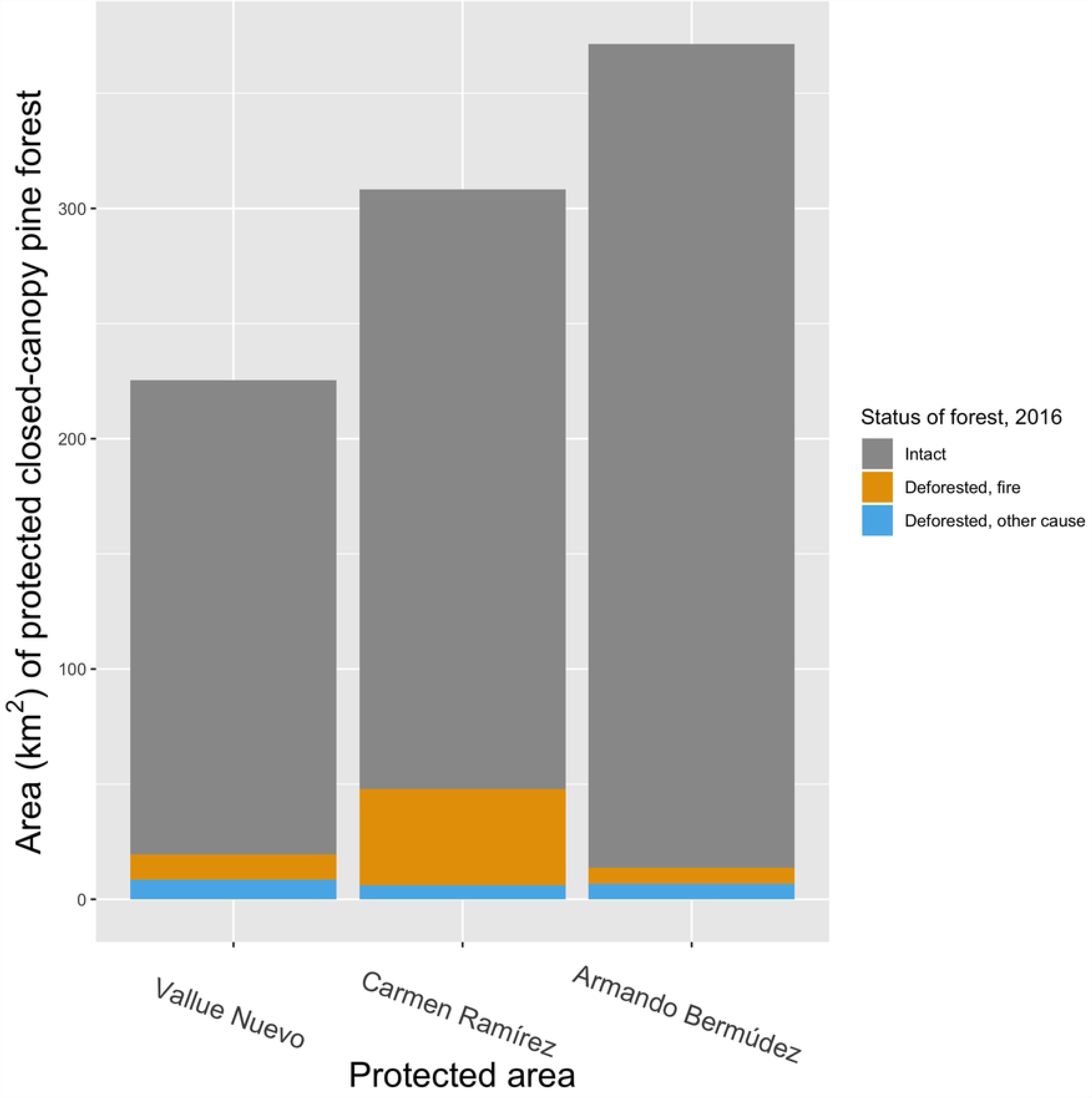
Area of closed-canopy Hispaniolan pine (*Pinus occidentalis*) in the Dominican Republic within protected areas that remained intact or was deforested (due to fire or other causes) between 2000 and 2016. Forest loss in the three most important protected areas - all classified as national parks (IUCN Category II) and collectively accounting for 67% of the total protected area for this forest type - varied due to the higher losses from wildfire in José del Carmen Ramírez National Park. Loss of forest cover from other sources was similar across the three protected areas.

**Fig 4.**
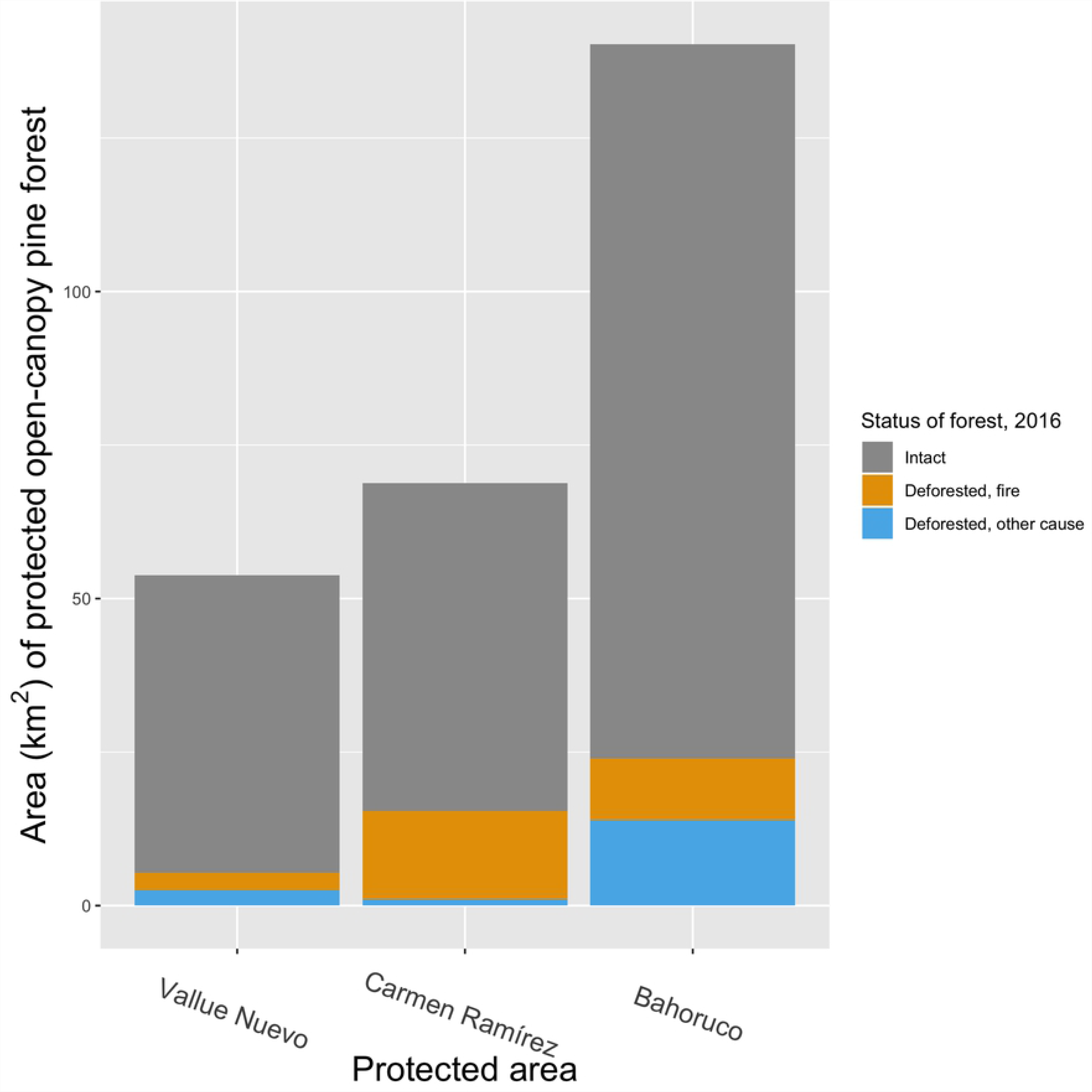
Area of open-canopy Hispaniolan pine (*Pinus occidentalis*) in the Dominican Republic within protected areas that remained intact or was deforested (due to fire or other causes) between 2000 and 2016. Forest loss in the three most important protected areas - all classified as national parks (IUCN Category II) and collectively accounting for 72% of the total protected area for this forest type - varied due to the higher losses from wildfire in José del Carmen Ramírez National Park and to deforestation from causes other than fire in Sierra de Bahoruco National Park.

Cloud forest losses within protected areas ranged from <1 km^2^ in Armando Bermúdez National Park to 11 km^2^ in JC Ramírez, or 17% of that park’s extant cloud forest (Fig. 5; S1 File). Roughly half (51%) of the cloud forest lost in JC Ramírez was due to the same wildfires that burned through the park’s pine forests. Other parks experiencing substantial loss of cloud forest were Valle Nuevo National Park, which lost 7.7 km^2^ (4.1% of its extent in 2000), and Bahoruco, which lost 7.6 km^2^ (8.2%). Loss of cloud forest in these two parks was driven primarily by processes other than fire (only 26.3% and 8.8%, respectively, of the deforestation in each was caused by fire).

**Fig 5.**
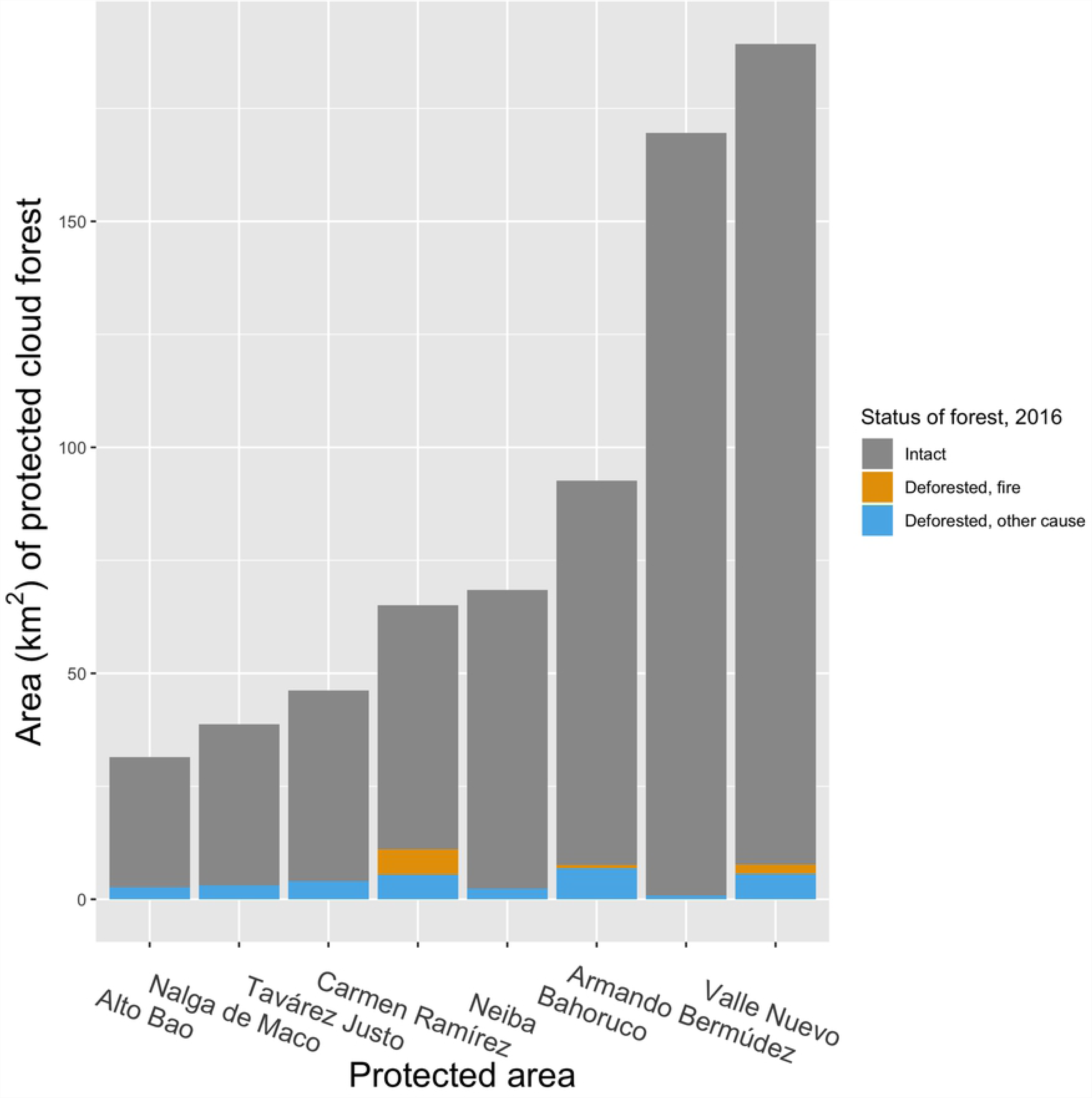
Area of cloud forest in the Dominican Republic within protected areas that remained intact or was deforested (due to fire or other causes) between 2000 and 2016. Forest loss in the eight most important protected areas - all classified as national parks (IUCN Category II) except for Alto Bao, a forest reserve (IUCN Category V), and collectively accounting for 94% of the total protected area for this forest type - was mostly due to sources other than fire. Fire was an important source of deforestation only in José del Carmen Ramírez National Park; Sierra de Bahoruco National Park and Valle Nuevo National Park both lost large areas of cloud forest from causes other than fire. Moist broadleaf forest losses were greatest in Los Haitises National Park, which lost 26 km^2^ (14.6%), almost all (94.2%) due to causes other than fire (Fig. 6; S1 File). Bahoruco experienced significant losses of moist broadleaf forests, too (8.6 km^2^, or 8.4% of the amount estimated to exist in 2000). Although fire was not generally an important cause of loss of this forest type, the 2005 fires that burned in JC Ramírez accounted for 71.4% of the observed moist broadleaf deforestation in that park, which totaled 4.1 km^2^.

**Fig 6.**
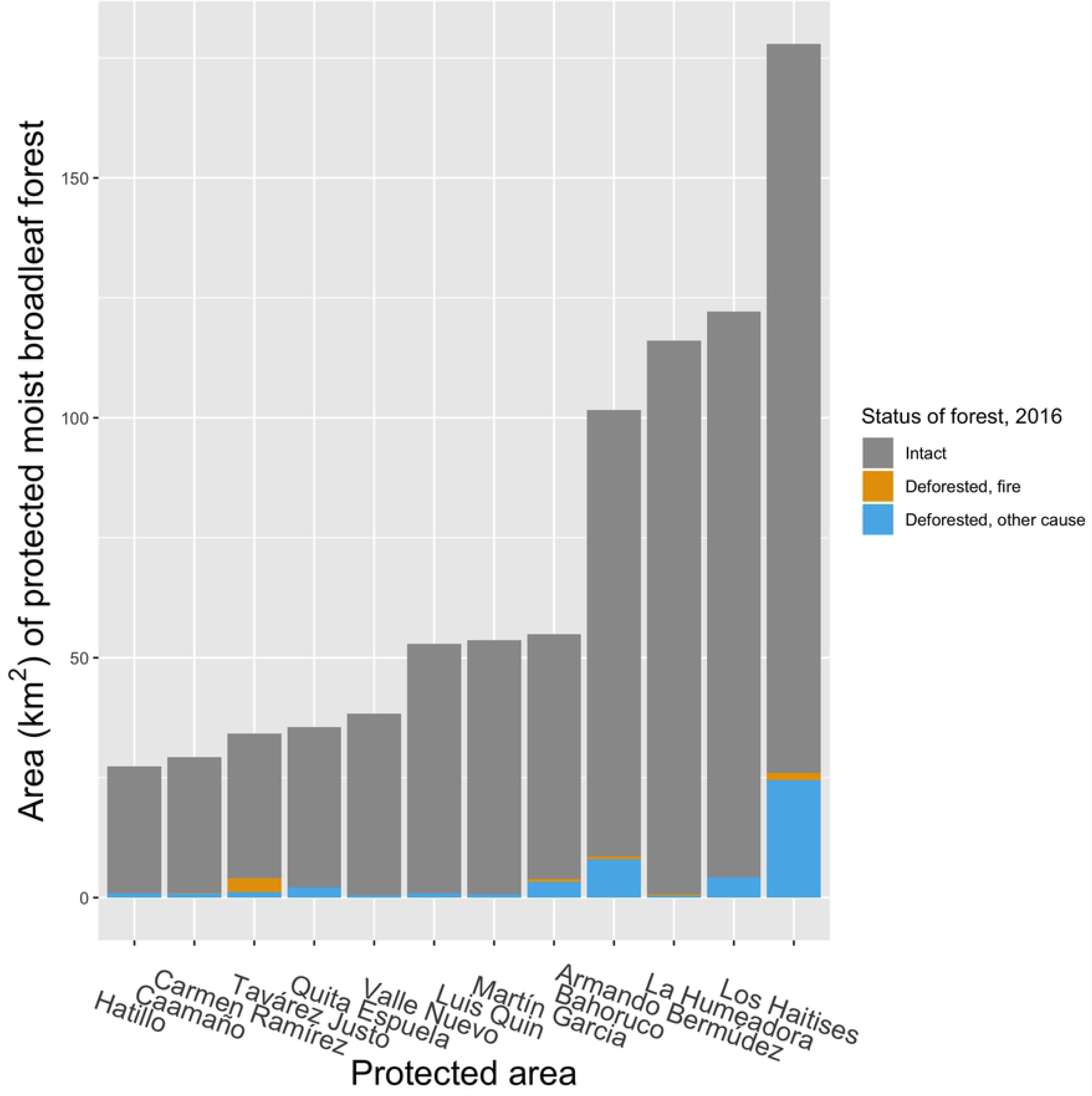
Area of moist broadleaf forest in the Dominican Republic within protected areas that remained intact or was deforested (due to fire or other causes) between 2000 and 2016. Forest loss in the twelve most important protected areas for moist broadleaf forest, which collectively account for 75% of the total protected area for this forest type, was concentrated in a single National Park (IUCN Category II), Los Haitises, and was due almost entirely to causes other than fire. Loss of semi-moist broadleaf forest was most pronounced in Bahoruco (13.9 km^2^, or 15.5% of the 2000 total extent) and Cotubanamá National Park (formerly Del Este National Park; 6 km^2^, or 2%; Fig. 7; S1 File). Bahoruco also led all parks in the amount of dry forest eliminated, with 7.8 km^2^ (5.1%) lost over the course of this study (Fig. 8; S1 File). Despite it relatively small size, Cerro Chacuey Natural Reserve was another noticeable hotspot of deforestation, losing 4.7 km^2^ or 35.2% of its extant dry forest (S1 File). Fire was unimportant as a driver of deforestation of both semi-moist broadleaf and dry forests in this study.

**Fig 7.**
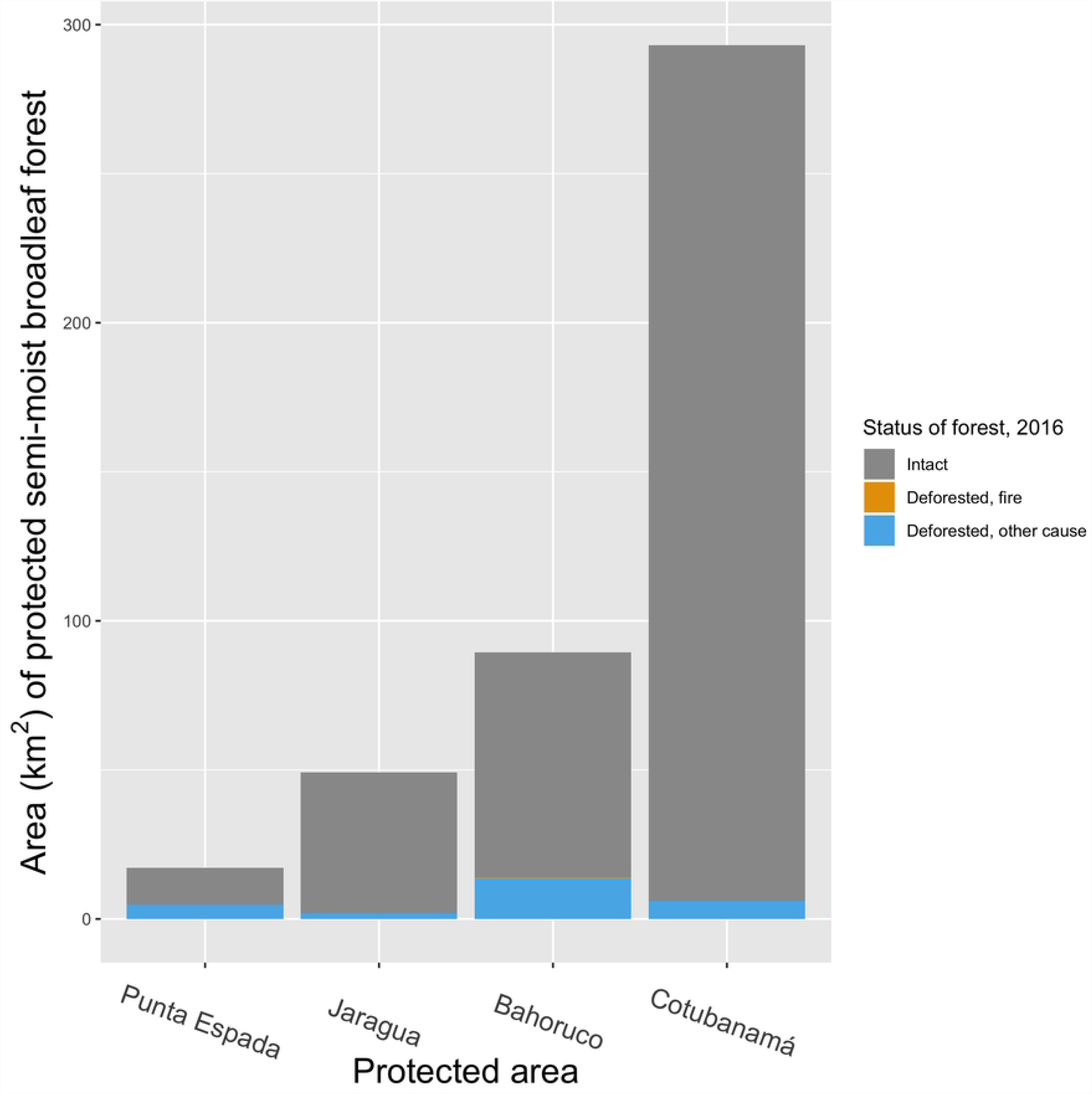
Area of semi-moist broadleaf forest in the Dominican Republic within protected areas that remained intact or was deforested (due to fire or other causes) between 2000 and 2016. Deforestation in semi-moist broadleaf forest was concentrated in two National Parks (IUCN Category II), Punta Espada and Sierra de Bahoruco. Collectively these four protected areas account for 75% of the total protected area for this forest type. Fire was an insignificant cause of deforestation in this forest type.

**Fig 8.**
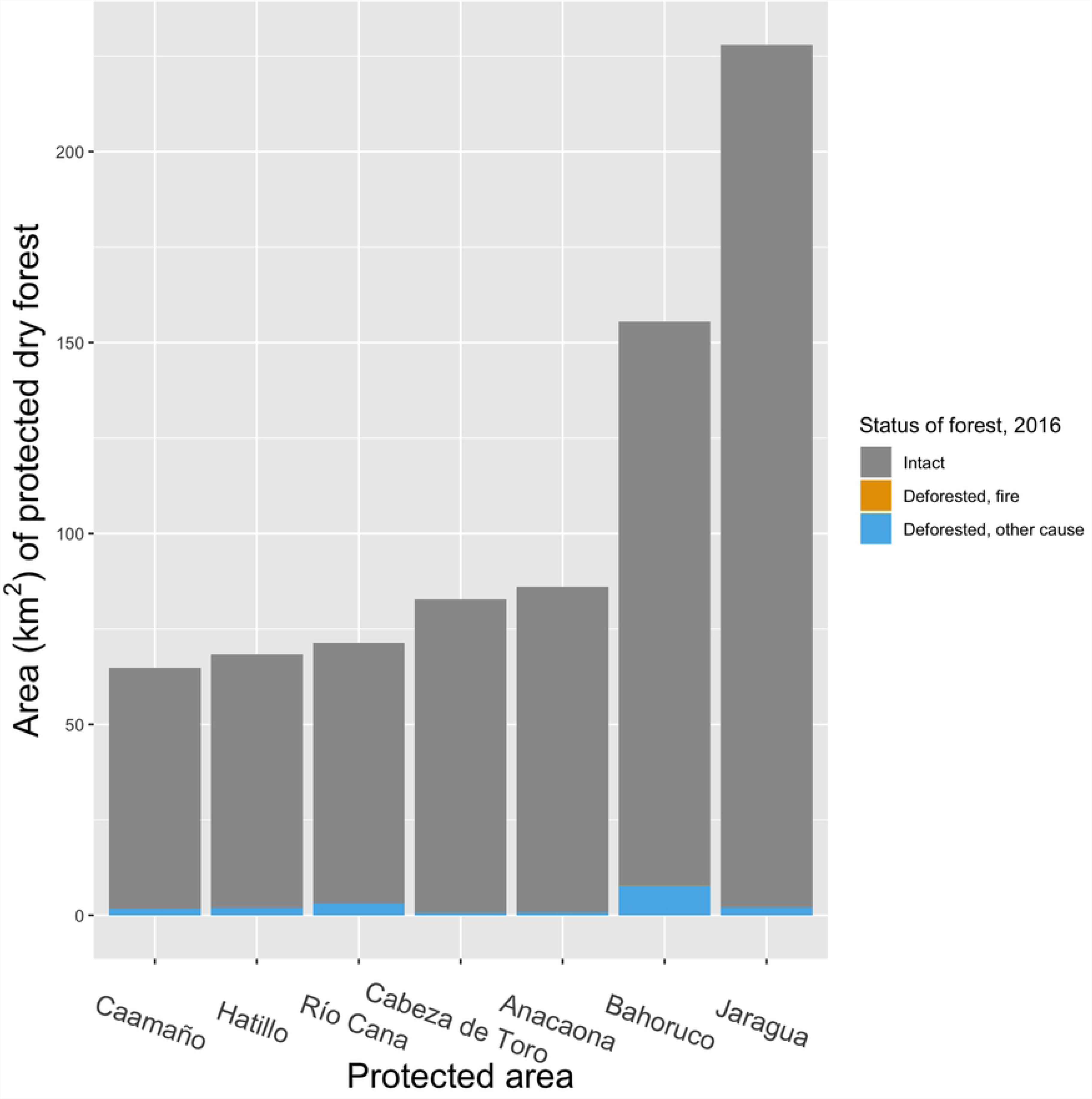
Area of dry forest in the Dominican Republic within protected areas that remained intact or was deforested (due to fire or other causes) between 2000 and 2016. Loss of protected dry forest was relatively high in Sierra de Bahoruco National Park. Collectively these seven protected areas account for 72% of the total protected area for this forest type. Fire was an insignificant cause of deforestation in dry forest.

## Discussion

Forest cover in the DR shrank substantially between 2000 and 2016, from nearly 45% to just under 40% of its territory, an overall decline of 11.1% and an annual deforestation rate of 0.7%. This rate was much higher than the 0.38% annual net rate of deforestation estimated for the tropics as a whole by Achard et al. [11] and for the mainland Neotropics in particular [27]. However, our estimated deforestation rate is comparable with the long-term 0.71% annual rate of forest loss estimated for the Amazon forest of Brazil [28].

Other recent studies, all using satellite sensor data, have reported qualitatively similar changes: Heino et al. [21] estimated that forest cover in the DR declined by 1,863 km^2^ between 2000 and 2012 (or 7% out of an estimated total area of 26,952 km^2^, a ∼0.6% annual decline), while Sangermano et al. [20], using a different approach, estimated a slightly slower rate of deforestation in the DR between 2000 and 2011, reporting a net loss of 518 km^2^ (or 5.3% of total forest area, a ∼ 0.5% annual decline). Furthermore, in keeping with our finding of widespread deforestation in all forest types and without respect to protected-area status, both Heino et al. [21] and Potapov et al. [10] reported significant declines in the extent of intact forest in the DR. In these studies, intact forest was defined as forest blocks > 500 km^2^ in area and minimally influenced by human activity, essentially all of which falls within the boundaries of protected areas in the DR. Potapov et al. [10] found a 29% decline in the extent of intact forest in the DR between 2000 and 2013, mostly due to losses caused by fire, whereas Heino et al. [21] estimated an 8.6% decline in intact forest.

In contrast to these trends, the Food and Agriculture Organization (FAO), using reports provided by the government of the DR, reported a 33.4% increase in forest cover between 2000 and 2015, an annual gain of 1.9% (FAO 2015). The methodology underlying this estimate is not identified in the FAO report, but it was probably derived from the national land-cover and vegetation maps produced by the Ministry of the Environment of the DR every few years. Elsewhere, national reports used by the FAO have been criticized as unreliable [29] and several studies have documented significant discrepancies between international estimates and nationally reported estimates of forest loss [9,30]. Furthermore, Romijn et al. [31] rated as low the capacity of the DR to carry out forest inventories and to monitor change in forest area, both of which are essential in generating reliable national reports on forest change. Given this, and given the consistency of estimates produced by international studies, we consider it unlikely that reforestation exceeded deforestation and instead have high confidence that the total area of forest in the DR declined from 2000-2016.

One possible source of error in our estimates of net deforestation is that our estimates of gain in the area of each forest type apply only to pixels falling within the mapped distribution of each forest type. Because we based our estimates of change in each forest type on its 1996 mapped distribution, we cannot rule out the possibility that areas categorized as another land-cover type in 1996 (e.g., subsistence agriculture) could have regrown into one of the forest types we analyzed. This would not have been captured by our analysis, thus leading us to underestimate forest gains during the period. However, the total gain in tree cover across all of the agricultural or otherwise anthropogenic land-cover types in the 1996 land-cover map was only 24 km^2^, so even if all of this gain reflected reversion to native forest cover, which is unlikely, it would account for only a small fraction of the 874 km^2^ of forest lost. Thus, we are confident that afforestation of agricultural or developed lands could not have materially affected our estimates of net loss.

As has been reported in other studies of deforestation in the Neotropics [30,32], we also found that deforestation in the DR tended to accelerate over time, with the exception of dry forest loss, which showed some evidence of a decline in the extent of deforestation after 2010. This slowing deforestation rate in dry forests could be because of the substitution of propane gas for wood charcoal – the main historical use of dry-forest trees – as the primary cooking fuel in the DR [33], a phenomenon also observed in Puerto Rico [34]. Charcoal trade went from roughly 1.6 million sacks in 1982 to just 49,000 in 2005, and its use as cooking fuel went from 90% of households in 1980 to just 10% in 2006 [35]. Although there are still some hotspots of illegal charcoal trade [36], especially in areas near the border with Haiti, quantifying its importance is difficult. Nonetheless, the widespread shift away from charcoal as the leading cooking fuel in the DR likely explains much of the observed drop in dry-forest deforestation rates.

### Loss by forest type and drivers

Outside of areas known to have burned, the data that we used do not provide direct insight into the drivers of forest loss. However, we can reasonably speculate that, with the exception of pine forests, the most likely cause for the observed forest loss is expanding agriculture. This is not only consistent with our field observations, but also in agreement with the findings from a comprehensive, national-level assessment which ranked agriculture as the leading cause of deforestation, accounting for 55% of forest loss in the DR [37]. In comparison, the same study attributed only 26% of deforestation to timber harvesting, firewood collection, and wood-charcoal production.

The important role of agriculture in forest clearing in two montane national parks has also been highlighted in recent reports by Wooding and Morales [38] for Nalga de Maco National Park and León et al. [39] for Sierra de Bahoruco National Park. Both studies describe the expansion of a similar commercial agricultural system, consisting of sharecropping in a shifting-agriculture system established between a landless Haitian farmer and a Dominican who claims land ownership. Sharing arrangements can vary, but usually the farmer keeps most of the crop, which is typically short-cycle crops. León et al. [39] also described the recent establishment of more permanent forest conversion in the form of avocados (*Persea americana*) grown for export, plantations of which have actively expanded inside Sierra de Bahoruco National Park since 2008. The problem of agriculture within protected areas is not limited to montane parks, however; a study on the drivers of deforestation in the low-elevation Los Haitises National Park also identified farming as the leading cause. In this case, deforestation was driven by increased exports of taro root (*Colocasia esculenta*), the leading crop inside the Park [40].

Fire was the leading cause of forest-cover decline in Hispaniolan pine forests. Pine trees and their associated understory plants are not only resilient to fire, but depend on it for seed dispersal and germination [41] and thus, absent any additional disturbance, burned pinelands will likely recover [42]. Of concern, however, is evidence of emerging changes in fire regime that may pose a long-term threat to these forests. Whereas lightning during dry seasons was probably the leading cause of fire ignition in the past, today human activities are. The DR’s National Fire Management Strategy has identified as the leading causes of forest fires, in order of importance: farming activities (especially land preparation for short-cycle crops), renewal of cattle grazing pastures, intentional fires in protest against authorities, and accidental fires caused by abandoned cooking fires from hunters and parrot poachers [43]. Furthermore, the strategy highlights a new and complex threat: the expansion of the invasive molasses grass (*Melinis minutiflora*), which is highly flammable and has already been implicated in forest fires [43]. Changes in the seasonality, frequency, or intensity of fire may negatively affect even relatively resilient pine forests, let alone broadleaf forests that are ill-adapted to fire.

Cloud forest also experienced substantial losses due to fire. However, unlike pine forest, it is far less resilient to fire. Not only is cloud forest exceedingly slow to recover after fire [44], but exposure to repeated fire can lead to its replacement by other forest types [42]. The fire-related losses of cloud forest that we documented, therefore, may be permanent. This is very concerning as these montane forests not only host most of the unique, threatened species on the island, but also intercept water from rain and clouds year-round (e.g., [45]), allowing lowland human communities to thrive even in extremely dry areas. The 2005 fires that produced most of the fire-related deforestation in pine and cloud forests were exacerbated by drought conditions brought about by an El Niño event in late 2004 [24]. The climate of the DR is expected to grow warmer and drier under most scenarios of climate change [46], raising the possibility that fire - probably historically unimportant as a driver of change in cloud forest [24] - may become a far more important threat to cloud forest in the future.

### The impact of protected areas

Protected areas lost less tree cover than did unprotected areas, as has been shown previously both in the DR [20] and in other parts of the world [47-50]. However, protected areas also varied in terms of the degree of protection they afforded. Three mountain-based National Parks in particular - Sierra de Bahoruco, José del Carmen Ramirez, and Valle Nuevo - exhibited consistently high rates of deforestation across multiple forest types. A fourth National Park, Los Haitises, located at sea level, had notably high rates of deforestation in its predominant forest type, moist broadleaf forest. These four areas all share a common problem: expanding agriculture or cattle ranching operations. These activities are feasible in these protected areas because they have a relatively humid climate, and in the case of the three mountain parks, milder temperatures, conditions which allow for profitable farming operations without expensive watering systems. This is consistent with a recent global study on protected areas under pressure, which estimated that 21% of land within protected areas in the DR faced intense human pressure [51]. The low management effectiveness in some of the DR’s protected areas was also highlighted by a report from Sánchez [52], in which he measured a number of key management variables for the leading 35 protected areas (out of a total of 118 areas at the time). Of these, only four obtained a satisfactory management score (above 75%). Of the remaining 31 areas that failed to receive a passing score, ten showed evidence of ongoing decline in management effectiveness during the course of the three-year study. The lack of basic management attributes such as clear knowledge of protected area boundaries and the existence of management plans drove most of these low scores. From our observations in the field, besides a limited capacity to enforce existing protected-area laws, political patronage, local power structures, and corruption also play a role in limiting the effectiveness of protected areas.

Our findings also suggest that protected areas were more effective in reducing deforestation at lower elevations, particularly in dry forest. However, this could be attributed to several factors besides protection status, including the shift away from wood charcoal as cooking fuel in the DR, as well as the limitations that local climatic conditions impose on the development of agriculture and cattle ranching. These activities are only possible in dry forest sites with abundant, nearby freshwater resources, and often only after sizeable investments in irrigation infrastructure. Financing such investments often requires land titles, which can be difficult to obtain in legally protected areas. This agrees with the findings of Joppa and Pfaff [48], who also found that protected areas appear more secure when established in areas not highly valued for extractive resource uses. The apparently greater effectiveness of protected areas in areas of dry forest in the DR may thus simply reflect the low profitability of exploiting the resources that they contain, in contrast to the relatively lucrative opportunities afforded by the export-oriented agriculture that can be carried out in protected areas with more suitable climatic conditions.

### Policy implications

Although not typically considered a hotspot of deforestation, rates of forest loss in the DR are higher than regional averages and show no sign of decelerating. Our results reveal ongoing deforestation across the country, especially in moist forest types that are more valuable for agricultural development. Protected areas offered only modest reductions in deforestation for most forest types, highlighting a general lack of management effectiveness. As nations continue to expand their protected-area systems, there is an urgent need to undertake objective assessments of their effectiveness in meeting their goals, especially those pertaining to forest conservation. Satellite images and forest-cover analysis platforms, such as Global Forest Watch, offer an inexpensive and objective way to achieve this.

Continued deforestation in the DR poses a risk to the flow of critical ecosystem services, especially the provision of water by upland forests to lowland human communities, including the major cities and agricultural regions. Ongoing deforestation will also threaten the achievement of a number of the DR’s sustainable development goals, as well as meeting its Intended Nationally Determined Contribution under the Paris Agreement within the United Nations Framework Convention on Climate Change (UNFCCC). Widespread forest loss will also hinder the DR’s commitments to halt biodiversity loss as a party to the Convention on Biological Diversity, by placing at greater risk many unique, globally threatened species that depend on the country’s forests.

Addressing deforestation will require a better understanding of its causes. Although fire is an important driver of loss of forest cover in Hispaniolan pine forest, and occasionally in adjacent cloud forest, the vast majority of deforestation is driven by clearing for agricultural production [53]. More research into the local drivers of deforestation, its key actors, and associated social dynamics are needed. Efforts to stem deforestation will almost certainly involve stricter limits on large-scale agricultural commodity production within protected areas and the development of alternative livelihood opportunities for those practicing shifting agriculture. Shifting agriculture is in great part enabled by customary systems of land tenure in many rural areas of the DR that persist despite contravening laws and policies established by the central government. The critical role of land tenure in reducing deforestation, particularly under the REDD+ (Reducing Emissions from Deforestation and Forest Degradation) mechanism of the UNFCCC has been highlighted by a growing number of studies around the world [e.g., 54,55]. Addressing these issues is not easy, but will be crucial for securing the future of forests in the DR and in many other countries facing similar development pressures.

## Supporting information

**S1 File. Change in extent of major upland forest types within protected areas in the Dominican Republic, 2000-2016.**

